# Phylogeography, codon usages, and DNA barcoding of lesser short nosed fruit bat (*Cynopterus brachyotis* Muller, 1838) populations in Indonesia

**DOI:** 10.1101/2021.03.10.434719

**Authors:** Andri Wibowo

## Abstract

Fruit bat (Pteropodidae) is one of the mammals that is common in environments and widely distributed from subtropical to tropical Asia. Whereas the information on phylogeography of fruit bat *Cynopterus brachyotis* is still limited. From this situation, this paper aims to assess the phylogeography, codon usage, and DNA barcoding of *C. brachyotis* populations in Indonesia. Phylogeography was developed based on 657 bp of the mtDNA COI gene for all bat individuals and Bayesian inference to construct the phylogeny tree. The *C. brachyotis* populations in Indonesia are different to the populations from the Asia’s continent. The results show that *C. brachyotis* populations in Indonesia were divided into 3 distinct clades. A putative geographical barrier, recent, and rapid range expansion in the Sunda lineage associated with changes in sea levels, possibly coupled with related ecological differences, may have driven population divergence, allopatric, and sympatric speciation. Codon usage and high frequency were also contributing to the dispersal of *C. brachyotis* forming a distinct population.

## INTRODUCTION

Fruit bat (Pteropodidae) is one of the mammals that is common in environments. Fruit bat is widely distributed from subtropical to tropical Asia. In ecosystem, fruit bats occupies landscape from low land to hilly landscape. Widespread of bat distribution has attracted attention and work on bat study in phylogeography context. This is also supported by the high differentiation at the mtDNA level. Several species that expresses high differentiation include Drosophila (Desalle et al. 1987), reptiles (Godinho et al. 2006), and bats (Castella et al. 2001). As a result, series of studies have previously assessed the population genetics and phylogeography of mammals using mtDNA. Together those studies have revealed information on dispersal, phylogeography, population history, and genetic structure of studied mammals.

Indonesia is one of tropical countries having high diversity of mammals including fruit bats. Whereas most recent studies focused at local or regional scales only and few have attempted to study the phylogeography of bats. This is alo important since Indonesia’s diversity is divided into west and east regional diversities. The west parts of Indonesia have diversity resembles to Asian continent diversity and Australian continent has influenced the east parts of Indonesia. Then by studying the phylogeography of particular fruit bat *Cynopterus brachyotis* using mtDNA, it can be seen the diversity patterns of Indonesia’s mammals.

## MATERIALS AND METHODS

### Study area

This study was conducted in 3 locations across Indonesia region and following diversity division pattern. First location was in the west of Java island, second was in the east of Kalimantan, and the last location is in the central Sulawesi (Figure 1). All locations were similar located in hilly landscapes that were occupied by fruit bats. In each location, 5 fruit bats were studied.

**Figure 1.**
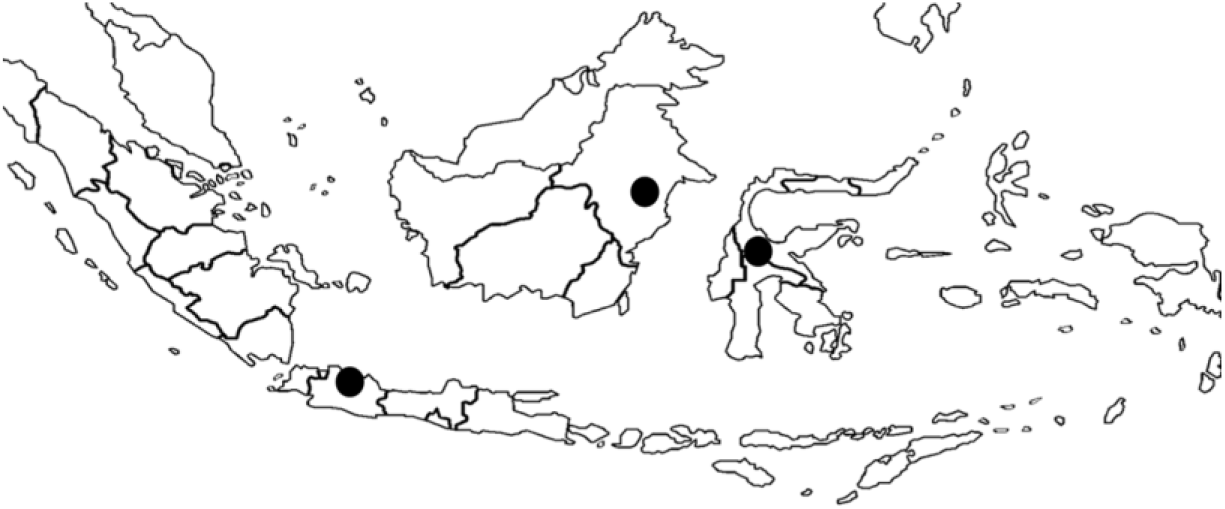
Studied locations (black dots) in west Java and east Kalimantan representing Asian diversity and central Sulawesi representing Australian diversity.

## Method

### DNA sampling

The DNA sampling was following methods of Neaves et al. (2016). Qiagen DNeasy blood and tissue kit following standard protocols or according to a high salt method were applied to extract genomic DNA. A base pair (bp) fragment of domain of the mtDNA was amplified using the specific primers. PCRs were carried out in 25 μl reactions using 100–500 ng of genomic DNA, 1 x reaction buffer, 2 pmol primers, and DNA polymerase. Thermocycling was performed with initial denaturation (94°C for 2 min), 38–45 cycles of denaturation (94°C for 20 s), annealing (60°C for 40 s), and extension (72°C for 50 s) followed by a final extension (5 min at 72°C). PCR products were cleaned and sequencing was resolved on a sequencer.

### Phylogeographic analyses based on mitochondrial sequences

The mitochondrial mtDNA sequence based phylogeographic analyses was following methods of Tu et al. (2017) and Bayesian inference based on Huelsenbeck & Ronquist (2001), Ronquist & Huelsenbeck (2003) and Ronquist et al. (2012). In a Bayesian analysis, inferences of phylogeny are based upon the posterior probabilities of phylogenetic trees. The posterior probability of the i_th_ phylogenetic tree (τi) conditional on an alignment of DNA sequences (X) can be calculated using Bayes theorem as follows:

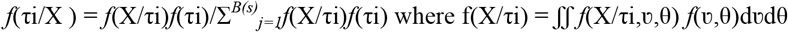

With the summation is over all *B(s)* trees that are possible for s species [*B(s)* = (2s-5)!/2^s-3^(s-3)! for unrooted trees and *B(s)* = (2s-3)!/2^s-2^(s-2)! for rooted trees], and integration is overall combinations of branch lengths (ν) and substitution parameters (θ). The prior for phylogenetic trees is *f*(τi) and is usually set to *f*(τi) = 1/ *B(s)*. The prior on branch lengths and substitution parameters is denoted as *f*(υ,θ). Typically, the likelihood function *f*(X/τi,υ,θ) is calculated under the assumption that substitutions occur according to a time–homogeneous Poisson process. The same models of DNA substitution used in maximum likelihood analyses can be used in a Bayesian analysis of phylogeny. The summation and integrals required in a Bayesian analysis cannot be evaluated analytically and it uses Markov chain Monte Carlo (MCMC) to approximate the posterior probabilities of trees. MCMC is a method for taking valid, albeit dependent, samples from the probability distribution of interest, in this case the posterior probabilities of phylogenetic trees.

## RESULTS AND DISCUSSION

The fruit bat populations in Indonesia are different to the populations from the Asia’s continent. This genetic variation can be observed in the form of distinct clades between Asia and Indonesia populations representing west Java, east Kalimantan, and central Sulawesi (Figure 2). Animals in Indonesia continents are originated from the Asia’s continent as this can be seen from water buffalo as an example. The current two distinct types (river and swamp) descended from different wild Asian water buffalo (*Bubalus arnee*) populations that are diverged some 900 kyr BP and then evolved in separate geographical regions. Swamp buffaloes dispersed through south east Asia and China as far as the Yangtze River valley (Zhang et al. 2020).

**Figure 2.**
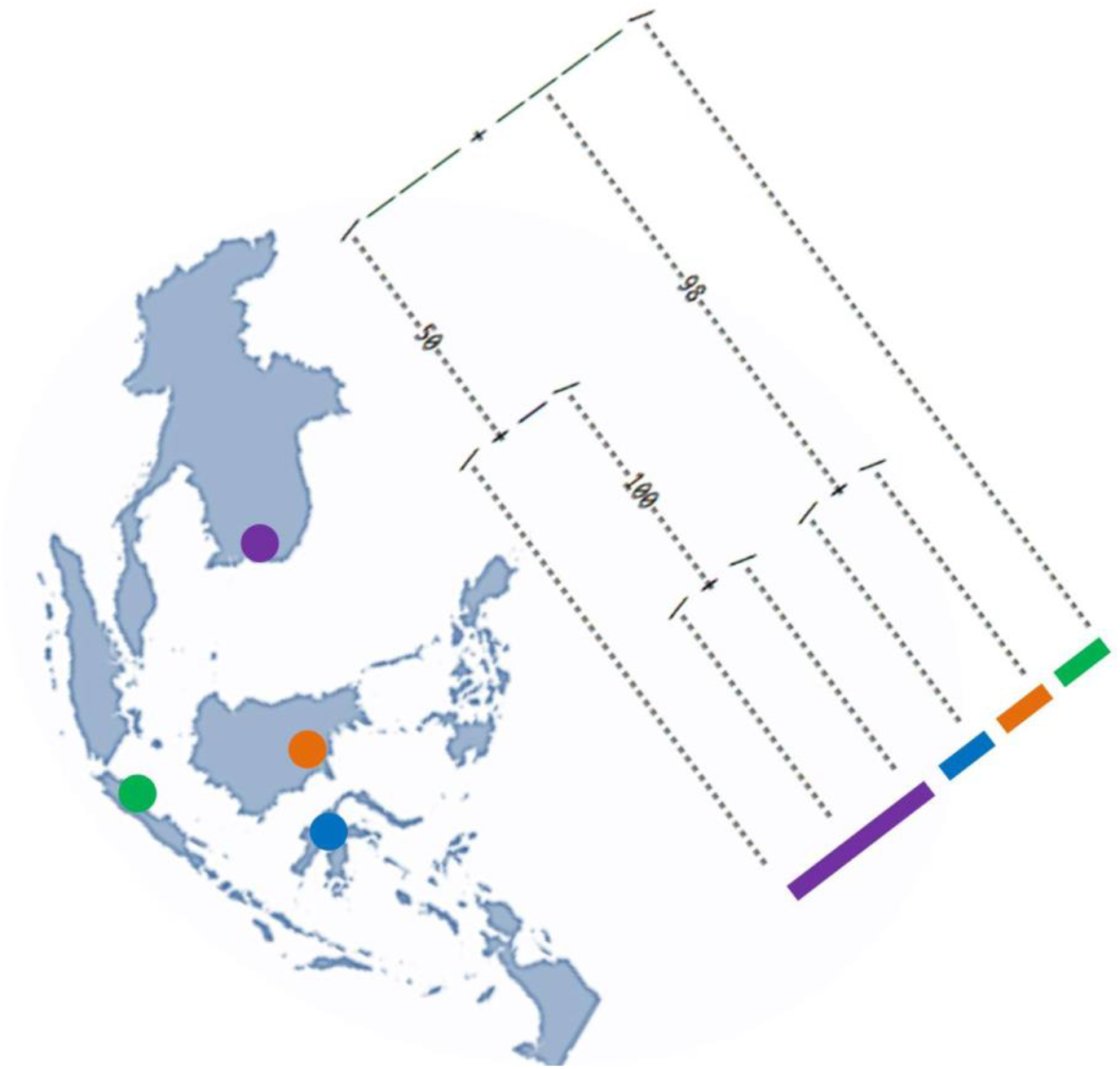
Phylogeography of lesser short nosed fruit bat (*Cynopterus brachyotis* Muller, 1838) populations in Asia continent and Indonesia (west Java, east Kalimantan, central Sulawesi).

Figure 3 presents a phylogenetic analysis of all lesser fruit bats samples with reliable geographical information (15 samples) using mitochondrialDNA revealed a large scale phylogeographic patterns across Indonesia. Those samples formed 3 distinct clades that very obvious for samples collected from west Java and central Sulawesi. Those locations were geographically separated representing distinct geographical regions and biomes whereas they have bats that share same ancestors. While east Kalimantan that geographically more closer to Java island, the bat population has different ancestors. The bats sampled from central Sulawesi were originated from a single ancestor and these reflect early divergences. The distinct population in central Sulawesi can be originated from a small bat population from west Java and east Kalimantan. While large populations were staying in east Kalimantan and forming distinct populations separated from west Java and central Sulawesi populations.

**Figure 3.**
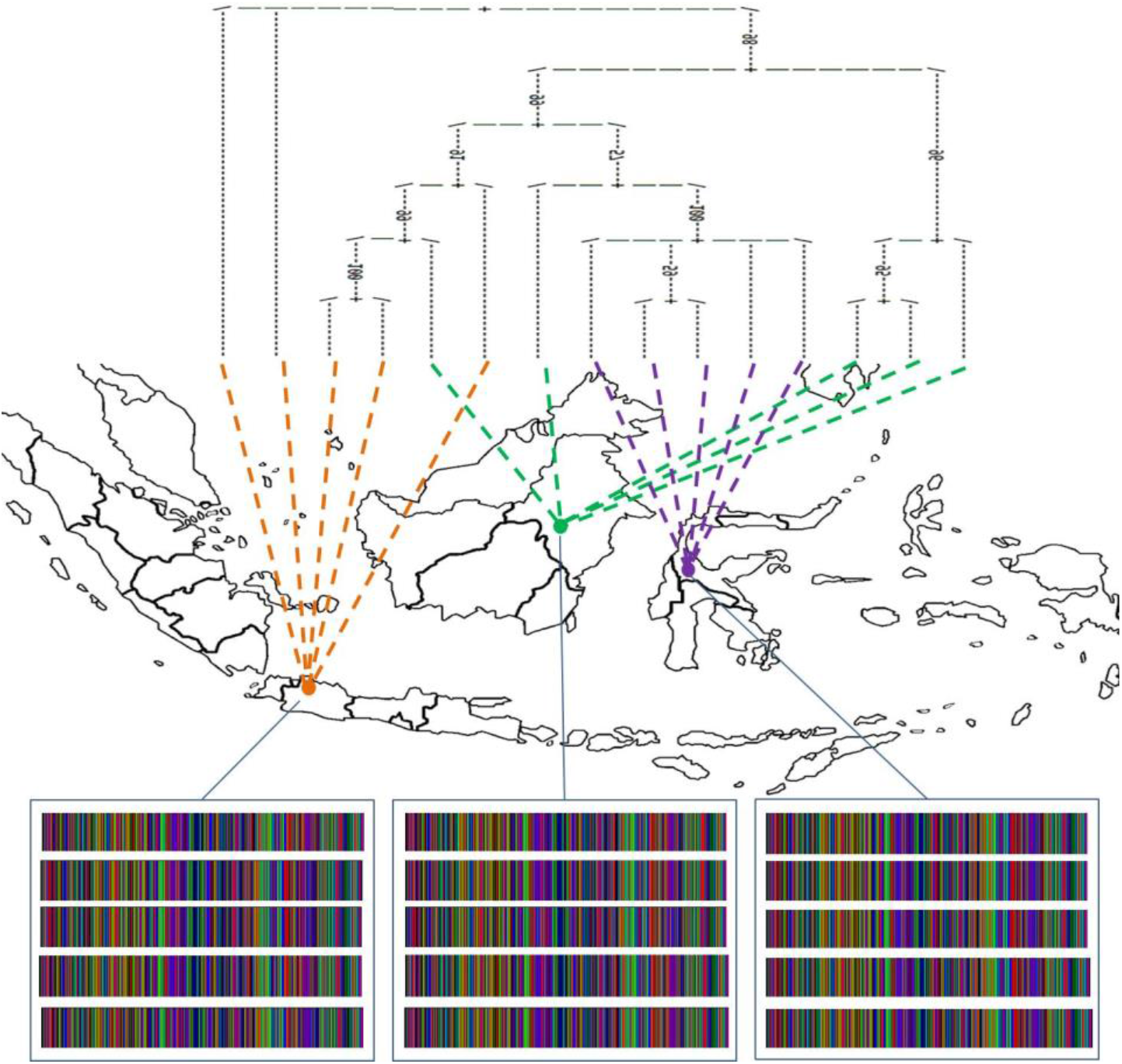
Phylogeography and DNA barcoding of lesser short nosed fruit bat (*Cynopterus brachyotis* Muller, 1838) populations in west Java, east Kalimantan, and central Sulawesi.

This research is important to complete the previous phylogeography research of fruit lesser bat as conducted by Campbell et al. (2004) that have identified six divergent mitochondrial lineages from India, Myanmar, Sulawesi, Sunda, and the Philippines with exclusion on Java and Kalimantan regions. In this study, there was a sympatric population with 2 bat populations from distinct ancestor living together in east Kalimantan. This finding is in corroboration with previous research that also reports sympatric populations of lesser fruit bats. A distinct population observed in west Java was related to a recent and rapid range expansion in the Sunda lineage and it was possibly associated with changes in sea levels during the Pleistocene. The geographic distribution patterns of the bat genetic diversity were also driven by population differentiation during the ice ages. Besides those factors, a distinguished genetic cluster representing distinct geographical areas can be related to the possible effects of past climate and geographic barriers to migration (Lompo et al. 2018).

In contrast to sympatric population, an allopatric population has been identified in this study since there are 2 distinct clades occupy west Java and central Sulawesi that shares same ancestor. This finding is comparable to allopatry of greater horseshoe bat, *Rhinolophus ferrumequinum* in southeastern Europe and Anatolia. Allopatric speciation of this species was related to suture zone in central Anatolia. Similar to those recorded in other animal species, showing the presence of more than one refugium within this region. The time of the split of these lineages that diverged in allopatry was dated to the Pleistocene. Natural and several putative biogeographic barriers can be potential factors limiting the distribution of mammals.

In this study, a population in central Sulawesi was having similar ancestor with another population originated from east Kalimantan. Those locations were separated by water with distance of 400 km and possible those areas were united during Pleistocene. A genetic variation and distribution of bat can occur in several locations that are separated. In South America, the populations of vampire bat (*Desmodus rotundus*) can be separated even within the distance greater than 1000 km. The bat populations in central Sulawesi were probably originated from the east Kalimantan populations and this was supported by the ability of bat to fly at far distance. This is explained by the distribution of vampire bat within a distance over than 1000 km since bat is able to fly 20 km from roost to feeding site in a single night (Martins et al. 2009). Bat can inhabit all the biomes that exist in the Neotropics, from seasonally flooded forests to semi-arid environments. This confirms that there is no identifiable physical barriers for dispersal and gene flow in the distributional range of bat in east Kalimantan reaching central Sulawesi.

This work describes the west Java, central Sulawesi, and east Kalimantan as a composite area for fruit bat with eastern and western components. The longitudinal division of this area has been recognized using parsimonious analysis of endemicity in vegetation, fishes, amphibians, reptiles, birds, and mammals. More recently, phylogeographic studies that used mtDNA described this structure in organisms as diverse as the most wildlife. All the studies that implemented time estimates yielded Pleistocene divergence times for this event (Kusmana & Hikmat 2015).

This study revealed at least there are 2 distinct populations in west Java and in east Kalimantan, while populations in central Sulawesi were clustered together with west Java population. The similar findings of distinct bat population have been observed by Coraman et al. (2013). According to their findings, among 12 species and the large *Myotis* complex, there were a total of 15 genetically distinct populations found and it represents biologically distinct taxa. Comparing the phylogeographic patterns of different taxa indicatse that distinct populations harbor genetically divergent populations and they should have higher priority in conservation practices. Considering a location contains richest bat fauna that is genetically distinct, protecting this particular distinct population is critically important for preserving the genetic diversity of the bats. Both regional and large-scale conservation strategies, which incorporate the distribution of genetic diversity, should be assessed and further ecological studies are needed to clarify the taxonomic relations of the identified clades.

Codon usage has been extensively used to assess an organism based on its genetically expressions (Mohajeri et al. 2016) and including comparisons among vertebrates (Kattoor et al. 2015) and mammals (Jørgensen et al. 2005). In this study (Table 1), bat populations from east Kalimantan were identified having the lowest codon frequencies and in contrast the central Sulawesi has the highest codon frequency among others. Codon usage and frequency are important and one of the determinant factors influencing adaptations of organism in even harsh environment, including gene expression to adapt drought (Mohasses et al. 2020). The central Sulawesi clade was separated from other populations and this Sulawesi population was having the highest codon frequency. For aerial animals, high codon frequency determines the ability to fly, disperse, and inhabit a new area. This explains the ability of bat originated from east Kalimantan to disperse in central Sulawesi and forming a distinct clade.

**Table 1.**
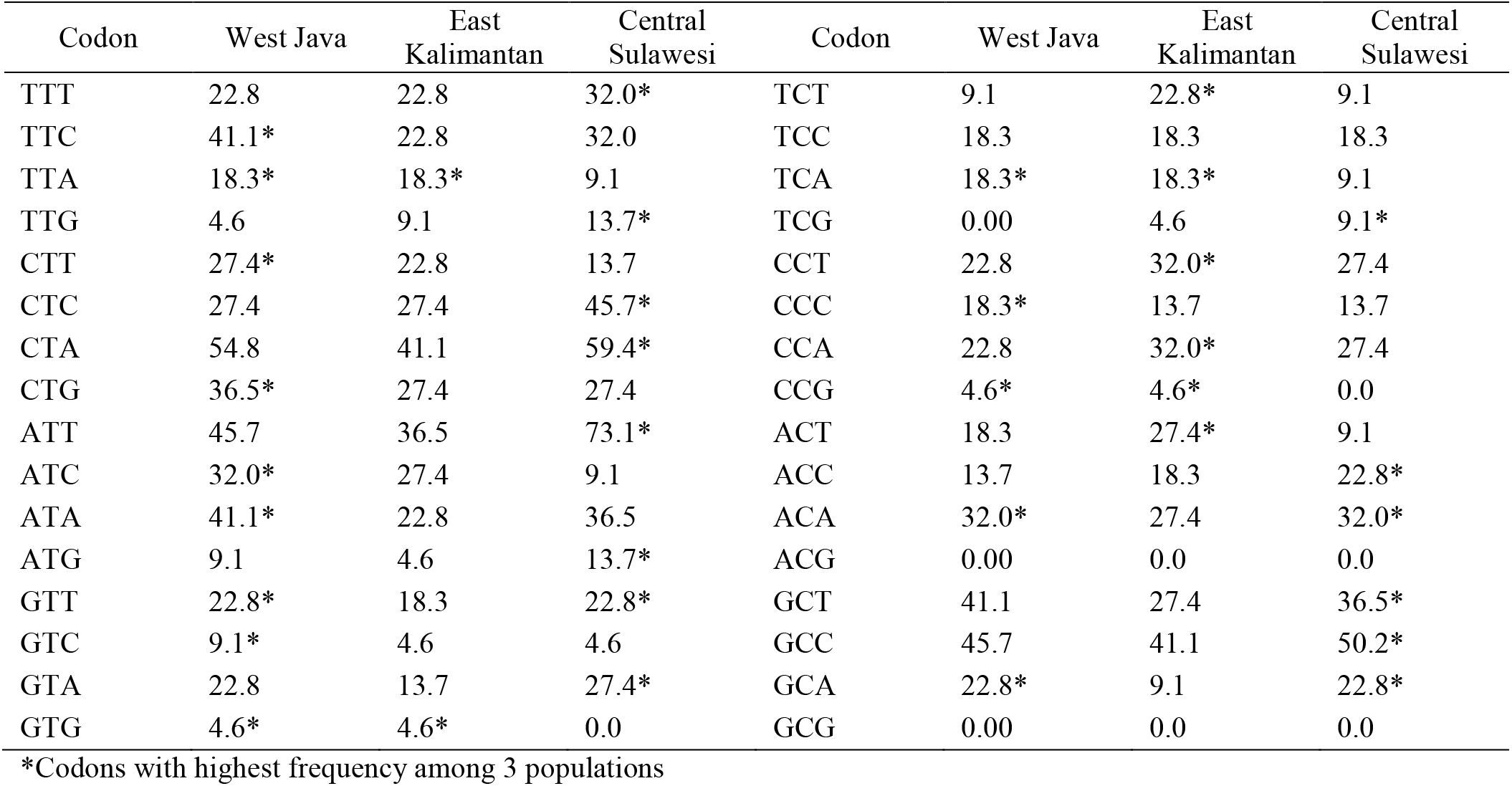
Comparison of codon usage in frequency among lesser short nosed fruit bat (*Cynopterus brachyotis* Muller,1838) populations in west Java, east Kalimantan, and central Sulawesi

| Codon | West Java | East Kalimantan | Central Sulawesi | Codon | West Java | East Kalimantan | Central Sulawesi |
| --- | --- | --- | --- | --- | --- | --- | --- |
| TTT | 22.8 | 22.8 | 32.0* | TCT | 9.1 | 22.8* | 9.1 |
| TTC | 41.1* | 22.8 | 32.0 | TCC | 18.3 | 18.3 | 18.3 |
| TTA | 18.3* | 18.3* | 9.1 | TCA | 18.3* | 18.3* | 9.1 |
| TTG | 4.6 | 9.1 | 13.7* | TCG | 0.00 | 4.6 | 9.1* |
| CTT | 27.4* | 22.8 | 13.7 | CCT | 22.8 | 32.0* | 27.4 |
| CTC | 27.4 | 27.4 | 45.7* | CCC | 18.3* | 13.7 | 13.7 |
| CTA | 54.8 | 41.1 | 59.4* | CCA | 22.8 | 32.0* | 27.4 |
| CTG | 36.5* | 27.4 | 27.4 | CCG | 4.6* | 4.6* | 0.0 |
| ATT | 45.7 | 36.5 | 73.1* | ACT | 18.3 | 27.4* | 9.1 |
| ATC | 32.0* | 27.4 | 9.1 | ACC | 13.7 | 18.3 | 22.8* |
| ATA | 41.1* | 22.8 | 36.5 | ACA | 32.0* | 27.4 | 32.0* |
| ATG | 9.1 | 4.6 | 13.7* | ACG | 0.00 | 0.0 | 0.0 |
| GTT | 22.8* | 18.3 | 22.8* | GCT | 41.1 | 27.4 | 36.5* |
| GTC | 9.1* | 4.6 | 4.6 | GCC | 45.7 | 41.1 | 50.2* |
| GTA | 22.8 | 13.7 | 27.4* | GCA | 22.8* | 9.1 | 22.8* |
| GTG | 4.6* | 4.6* | 0.0 | GCG | 0.00 | 0.0 | 0.0 |

\*Codons with highest frequency among 3 populations

## Notes

### Competing Interest Statement

The authors have declared no competing interest.

